# Harmonized decay classification for dead wood in Nordic national forest inventories

**DOI:** 10.1101/2023.09.04.556206

**Authors:** Tuomas Aakala, Juha Heikkinen

## Abstract

Dead wood quality is recorded as a biodiversity indicator and in estimating forest ecosystem carbon storage, using decay classification systems. In large-scale national forest inventories (NFIs), these systems are typically slightly different among countries, but harmonizing them would allows analyses over much broader scales and the use of larger data sets. Here, we developed a harmonized decay classification for the NFIs of Finland, Norway, and Sweden, using wood density as the harmonizing criterion. We sampled 441 pine, spruce, birch and aspen in different conditions and decay classes, measured their density and developed the harmonized classification for each species and dead wood type (standing and fallen dead wood). The assignments relied on minimizing within-class variance in wood density in the harmonized classes. Assigned into three (standing dead wood) and four (fallen dead wood) classes, the harmonization led to some losses of information, especially in the advanced decay stages in Finnish and Norwegian classifications. Importantly, the harmonized classes included slightly different decay classes from the national classifications, depending on the tree species and type. This is an advantage over the expert assessment that typically treat all species and types of dead wood the same way despite clear differences in decomposition pathways.

## Introduction

Dead wood is considered a key biodiversity indicator and plays an important role in forest carbon stocks and fluxes (Russell et al. 2015, Parajuli and Markwith 2023). It is hence tracked as an important variable in biodiversity assessments and quantified in carbon accounting. These information needs are likely to grow in importance in the future with efforts to halt and reverse the decline of biodiversity and with changing role of dead wood in carbon fluxes in a warmer world.

In many countries, dead wood is quantified as part of the national forest inventory programs (NFIs; Woodall et al. 2009). In Finland, Norway, and Sweden (henceforth: Fennoscandia), coarse recently dead or hard dead wood fraction of the total dead wood pool has been measured since the onset of the NFI programs 100 years ago. However, the entire coarse dead wood pool (limited to pieces larger than certain minimum size) has been incorporated in the measurements only since the 1990s, originally included primarily for biodiversity conservation reasons.

Upon the inclusion of dead wood in NFIs, their quality has been assessed as decay classes in all three countries, based on a combination of visual and tactile criteria (Storaunet and Rolstad 2015, Jonsson et al. 2016, Korhonen et al. 2021). These classifications are indicative of the habitat quality of the dead wood piece, but they are also a useful predictor for the wood density and the amount of carbon it still holds (Sandström et al. 2007). However, each Fennoscandian country has its own classification system for this purpose.

NFI programs produce powerful tools for analysis of forest structures and their changes. Their usefulness can be further increased by compiling combined data sets over NFIs of different countries, but this often requires harmonization for many of the variables (Stokland 2003, Rondeux et al. 2012, Ståhl et al. 2012). Such combinations may then encompass broader climatic and edaphic gradients, more variation in species and species communities, and different administrative units with differing forest uses and policies, and providing comparable data over larger regions (Gschwantner et al. 2019). As long-term data sets greatly increase in value with accruing re-measurements, harmonizations by assigning existing variables to harmonized variables (using so-called bridges, Ståhl et al. 2012) are obviously preferable over restarting measurement series with new classifications.

We present here a quantitative approach to harmonizing dead wood decay classes, as alternative to earlier approaches to harmonization relying on expert opinion (Stokland 2003, Rondeux et al. 2012). Our approach employs decay class-specific wood density (the basis for biomass expansion factors) as the harmonizing criterion. As initial wood densities and decomposition pathways may differ among species and between standing and fallen dead trees, assigning national classes to harmonized classifications may not follow the same scheme for all species and/or dead wood types. Thus, we developed separate harmonizations for standing and fallen dead trees of the main tree species in boreal Fennoscandia.

## Material and methods

### Fennoscandian tree species

The forests of Finland, Norway, and Sweden are mostly within the boreal zone, with the southernmost parts belonging to the hemiboreal and temperate zones. The forests are dominated mostly by Scots pine (*Pinus sylvestris*) and Norway spruce (*Picea abies*), with two birch species (*Betula pendula* and *Betula pubescens*) as the most important deciduous trees, along with trembling aspen (*Populus tremula*), and a number of co-dominants including two species of alder (*Alnus glutinosa* and *Alnus incana*), rowan (*Sorbus aucuparia*), and goat willow (*Salix caprea*). In the hemiboreal and the temperate parts, deciduous trees include oak (Quercus robur), maple (*Acer platanoides*), linden (*Tilia cordata*), ash (*Fraxinus sylvatica*), and beech (*Fagus sylvatica*). Here, we focus on Scots pine, Norway spruce, the two birch species (pooled together) and aspen, which are the most abundant species throughout the region.

### Dead wood and decay classifications in the Fennoscandian NFIs

In the Finnish classification, standing dead trees (snags) and fallen trees (logs) or pieces of fallen trees are classified into four and five classes, respectively. Although there are small differences between snag and log classifications, for both of them the main criterion is the depth to which a knife firmly pressed into the wood penetrates (Korhonen et al. 2021). In decay class 1 (the most recent), knife penetrates only a few mm, in decay class 2, the wood is still firm but starts to soften so that knife penetrates 1-2 cm. In decay class 3, the wood is fairly soft and the knife penetrates 3-5 cm. In decay class 4, the stem wood is soft and knife penetrates all the way. Wood material of logs in decay class five is highly decomposed and can be pulled apart by hand. These logs are often fully covered by epiphytes, mostly those typical to forest floor. Stems are thus often visible as an outline on the forest floor. In addition to this main criterion, there is a set of additional criteria, related to presence/absence of bark, branches, stem shape or epiphytic species to aid the classification.

Similar to the Finnish version, Tthe Norwegian classification (Storaunet and Rolstad 2015) has four and five decay classes for snags and logs, respectively. Dead trees in decay class 1 have died recently, have their bark still intact or the bark is recently loose due to bark beetle attacks. In the second class, bark is loose with well-developed mycelia between bark and wood. The wood is starting to decompose in the outer stem, with knife penetrating up to 3 cm into the wood. In decay class 3, outer parts of the stem wood are decomposed, and wood can be broken apart with a knife. The inner core of the stem is still hard. In decay class 4, wood is decayed throughout, with a loss of shape and absence of a hard inner core. Stem is often overgrown. In decay class 5, only fragmented parts and contours of completely decayed logs remain.

The Swedish decay classification (Jonsson et al. 2016) starts from class 0 that includes recently dead trees with green leaves or needles and/or fresh phloem. In decay class 1, over 90% of the stem volume consists of hard wood with a hard surface, very little influenced by decomposers. In decay class 2, trees are slightly decayed, with 10-25% of stem volume soft with the remaining part hard dead wood. Sharp metal objects (in practice, a knife) can be pushed through the outer part but not through the entire sapwood. In decay class 3, 26-75% of stem volume is soft or very soft wood. In decay class 4, trees are highly decayed, with 76-100% of stem volume soft or very soft. Although sharp objects can be pressed through the entire stem, hard inner cores may also be encountered.

### Data

Following a hands-on training with NFI field crews in Finland, Norway and Sweden in 2018 by the first author, we sampled a total of 441 dead trees in different parts of Finland. As within-species wood density is known to vary due to a number of different reasons (e.g., tree growth rates, latitude), we sampled trees in different growing conditions (latitudinal gradient) and growth histories (both managed and protected forests). For this, a 500 x 500 m grid of points were overlain on conservation areas in Finland, and 12 locations of these were chosen for sampling (Fig. S1 in Supplementary material). Sampling effort was divided so that the samples included areas from southern, middle, and northern boreal vegetation zones. Samples were then obtained from the selected conservation areas, and (for logistic reasons) from managed forests in the vicinity of these areas.

At each location, we searched for dead wood of different species, position (standing or fallen), and decay class within 500 m of the sample point. This opportunistic sampling strategy was used because finding samples in rare decay classes would have been prohibitively time consuming if it was based on a systematic or a random sampling procedure within the stands. We assumed that randomizing the location was sufficient to capture the range of wood densities in each decay class, position, and tree species. This approach was chosen with the aim of sampling a wide range of material for the decay class harmonization, but it should be noted that it is unlikely to produce unbiased, national-level estimates the way outlined in Sandström et al. (2007), or as discussed by Russell et al. (2015).

Once we located a tree, we classified it into decay classes, using all three national classifications. Two samples were extracted per tree, one at 25% and another at 75% of the length of the part of the stem that was over 10 cm in diameter (the minimum diameter in Finnish and Swedish dead wood measurements). Each sample was a disc, cut with chainsaw. Sample fresh volume was computed as the volume of a cylinder, based on two measurements of outer diameter at 90-degree angles to one another, and two thickness measurements as done in the field. Aspen snags in conservation areas were sampled differently by cutting out partial cross-sections so that a volumetric sample was obtained without felling the snag (as per the terms of the research permit for conservation areas). For trees in advanced decay stage covered with vegetation, the vegetation on top was removed prior to cutting. Highly decomposed logs were first cut with a chainsaw and then scooped into a plastic bag. Dimensions were then measured from the stem on the ground from the cut surface. At each locality, a maximum of two trees per species and decay class and position (snag or log) were sampled.

Samples were transported to a lab and stored in a freezer, until dried to constant weight (for 2 weeks in 65°C) and weighed. Density of an individual tree was obtained as a weighted average of the two samples, computed as the dry mass divided by the field-measured volume. Weighting was based on the cross-sectional area of each sample. With this approach we aimed at measuring wood density as it is measured in the field in the NFIs, i.e., without accounting for hollows etc. that are sometimes accounted for when determining density.

### Statistical analysis

All three national classifications have five classes (in practice, four for snags). We clustered the samples into three (for snags) and four (for logs) classes. We then tested all possible combinations of national classifications in the harmonized classification. The combination of classes into harmonized classification that minimized within-class variance was considered the optimal way to assign classes in the national classification into the new classification. In practice, we used the within-cluster sum of squares as the criterion. The computations were done in R, using the fpc-package (Hennig 2023), and ggplot2 (Wickham 2016) for visualizations.

## Results

### Sample trees and wood densities in decay classes

We sampled a total of 441 trees. Average number of samples per decay class, species and position (Table 1), was 12.2 for the Finnish and Norwegian classes, and 11.0 for the Swedish classes. Aspen samples were fewer than for other species. Within-species, rare decay classes had a smaller number of samples, such as decay class 0 in the Swedish classification, and conifers in highly decayed snag classes, as these are rare in managed forests, and in the protected areas in southern Finland.

**Table 1.**
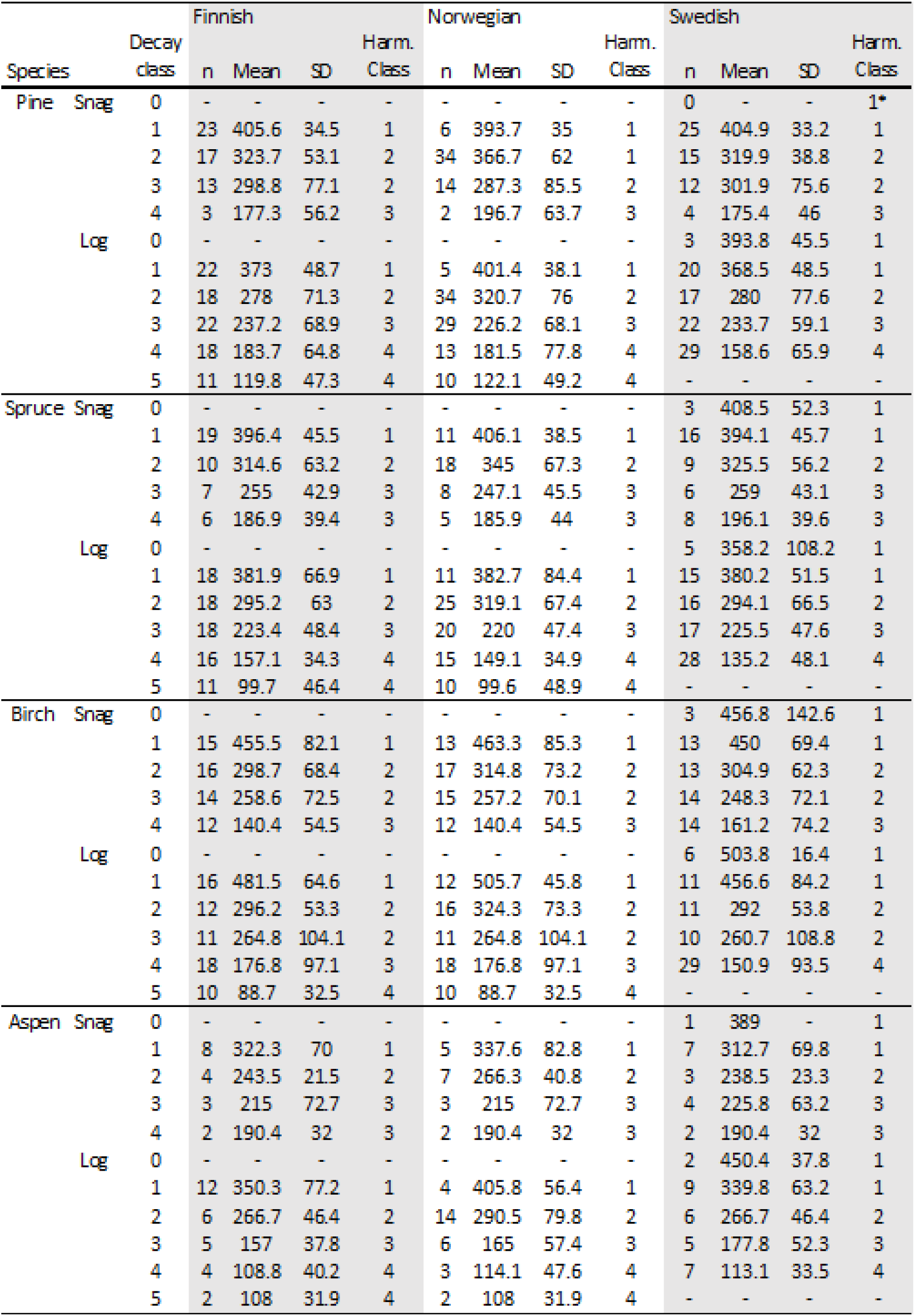
Means and standard deviations of wood densities in different decay classes for the three NFI classifications. Finnish and Norwegian classifications are from 1 to 5, Swedish classification from 0 to 4. Snags (standing dead trees) are not found in class 5 in the Finnish and Norwegian classifications.

In the national classifications, snag wood densities were similar in the two least decomposed classes (0 and 1) of the Swedish classification as in the least decomposed class in the Finnish and the Norwegian classifications (Table 1, Fig. S2a in Supplementary Material). In more advanced stages, mean wood densities for snags developed quite similarly across the national classifications, except that the Norwegian class 2 tended to have a slightly higher density than class 2 in the Finnish and Swedish systems.

For logs, the wood density followed a similar development as in snags, when considering the development in national classifications (Table 1, Fig. S2b in Supplementary Material). The main difference was that the most advanced decay class (class 5) in the Finnish and Norwegian classifications was clearly separated from earlier classes, and from the most advanced class in the Swedish classification (Swedish decay class 4).

### Harmonized classifications

Minimizing the within-group variance in selecting optimal groupings of decay classes based on wood density varied among species, and also within species so that the groupings were different for snags (standing dead trees) and logs (Fig. 1). The endpoints, most recent and the most advanced classes were systematically assigned into similar classes (Fig. 1). For the most recent classes, Finnish and Norwegian class 1 and Swedish classes 0 and 1 were always assigned to harmonized class 1. Logs in the most advanced class (class 5 in Finnish and Norwegian systems, and class 4 in the Swedish system), were assigned to harmonized class 4 in all cases. The mid-decay stages of the national classification systems showed more variation for both snags and logs as well as differences among species (Fig. 1).

**Fig. 1.**
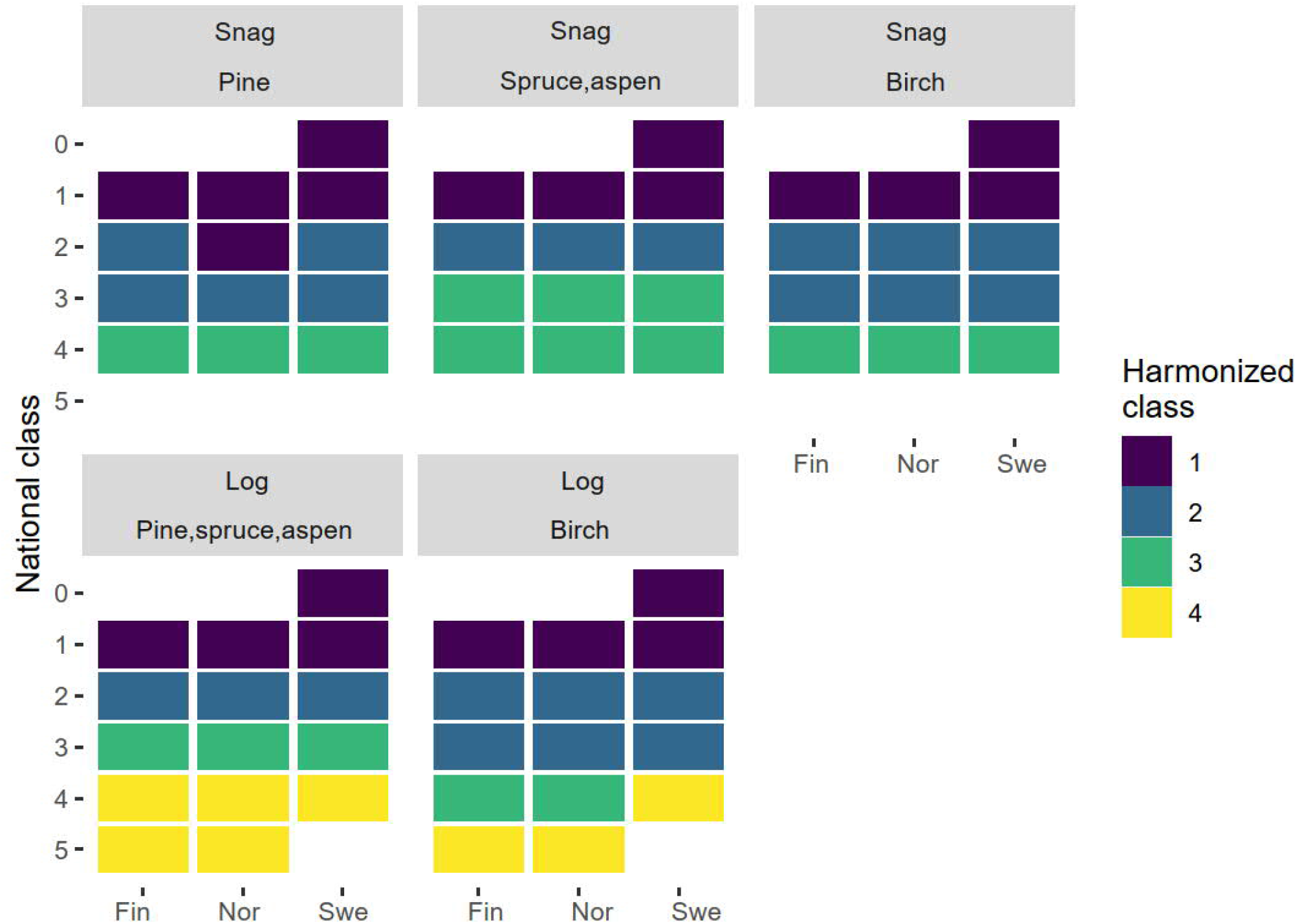
Schematic diagram of how trees in national classifications were assigned into the harmonized classes for different types of dead wood (snags, logs) and different species.

## Discussion

We presented here a harmonized decay classification for combining the Nordic National Forest Inventory dead wood data. Unlike earlier harmonizations (e.g., Stokland 2003), our approach was based on quantifiable criterion, wood density, instead of expert opinion. While in some of our cases expert opinions would likely have resulted in a similar outcome, the obvious advantage of this approach was that it accounts for differences in decomposition pathways. Here, this was visible as differences in the harmonization depending on the species and on whether the dead trees were standing or fallen.

The use of a reductive bridge (i.e., assigning 5-stage classifications to four classes) here led to modest loss of information from the perspective of wood density (and consequently carbon accounting), mainly in that for logs in the two most decomposed decay classes in the Finnish and the Norwegian classification were combined, even if they were well-separated in density in the original national classifications. In the Swedish classification, wood density in the two recent-most decay classes was very similar so that assigning Swedish classes 0 and 1 led to little loss of information on wood density. From the carbon accounting point-of-view, an improvement in the national classifications without losing information or comparability with earlier inventories would be an introduction of an additional decay class in the Swedish classification for the very late stages of decay (similar to decay class 5 in Finnish and Norwegian classifications).

It is noteworthy that the wood densities per decay class presented here are not necessarily an unbiased sample at the national level, because of the way sampling locations were selected geographically, and especially within each tree (Russell et al. 2015). Relative to the study of Sandström et al. (2007) where biomass expansion factors were developed for these same tree species and for the Swedish classification, our results were somewhat different: the average densities in Sandström et al. (2007) were consistently lower, especially in the early decay classes. This discrepancy may be due to a number of factors, but we suspect the main reason is the way sampling locations were selected within each piece of dead wood. Sandström et al. (2007) took three samples from different parts of the stem, one including the base of the stem that is often partly decomposed already while the tree is alive, potentially explaining the lower average densities. Other reasons may have been the subjective nature of the decay classification (Larjavaara and Müller-Landau 2010), and differences in the properties and growing conditions of the tree individuals that leads to differences in wood density at death. Nevertheless, it seems clear that a large sample size and a proper probability sampling for selecting the exact location where the sample is extracted along a stem would be preferable or even necessary for developing these estimates (Russell et al. 2015).

Given the proliferation of different decay classification systems in both research and practice (driven e.g., by the need to quantify carbon stocks and fluxes in forest ecosystems) a universal decay classification is unlikely to emerge with diverse criteria for assigning trees to decay classes (Russell et al. 2015). However, simple but objective approaches to harmonize, such as the one presented in this study would facilitate meta-analyses or allow pooling data over larger regions.

## Acknowledgements

We thank the field crews of the NFIs for the field training on the decay classifications. The work was funded by the Kone Foundation.

## Data availability

The sample data used in this work is openly available at DOI: 10.6084/m9.figshare.24081309

This dataset includes the data used here, and in addition a smaller number of samples for other boreal deciduous trees.

