## Supplementary material for "Harmonized decay classification for dead wood in Nordic national forest inventories"

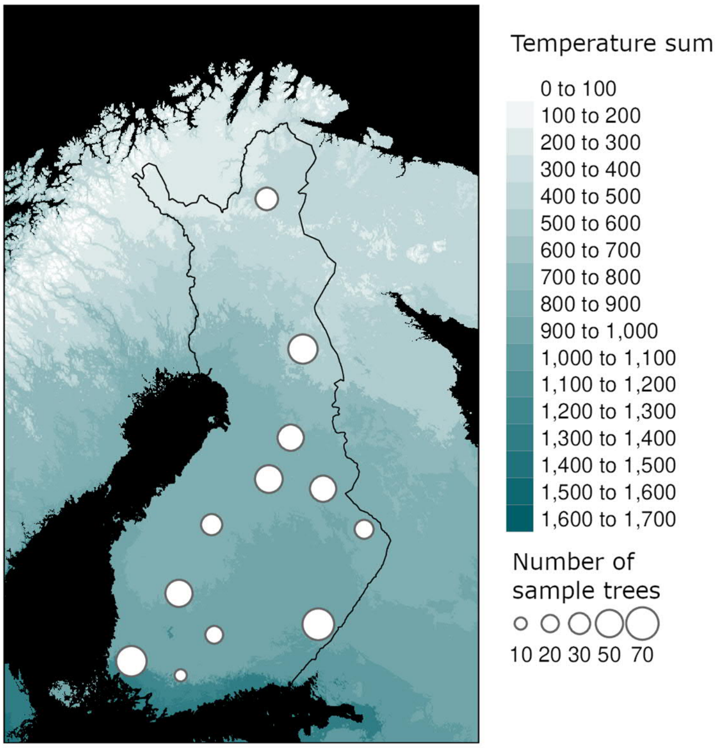


**Fig. S1.** Sampling locations on a map Finland showing the gradient of mean annual temperature. At each locality, dead trees were sampled within the protected area and/or state-owned managed forests close by.

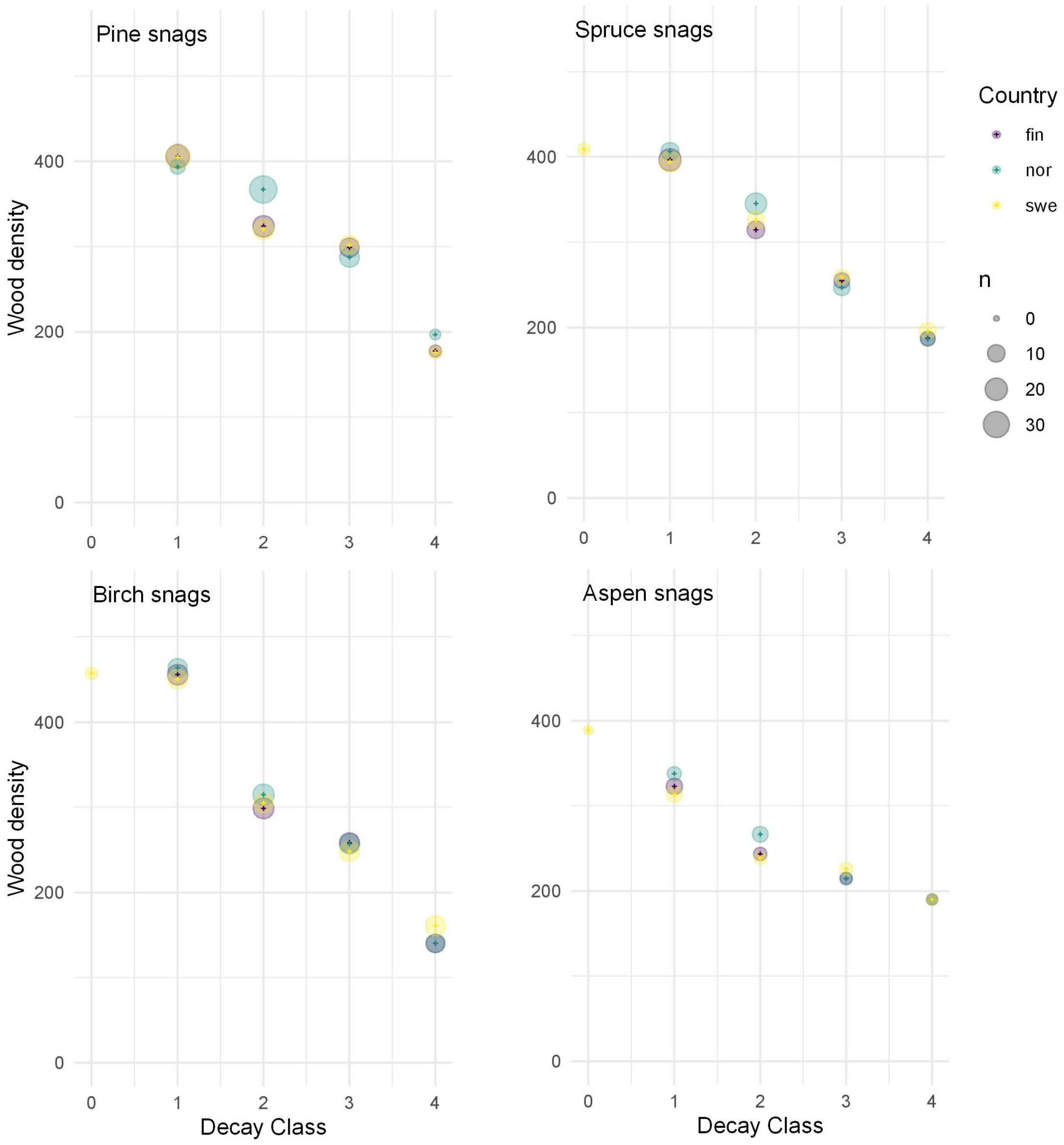


**Fig. S2a.** Average snag (standing dead tree) wood densities in the national classification systems of Finland (fin), Norway (nor), and Sweden (swe).

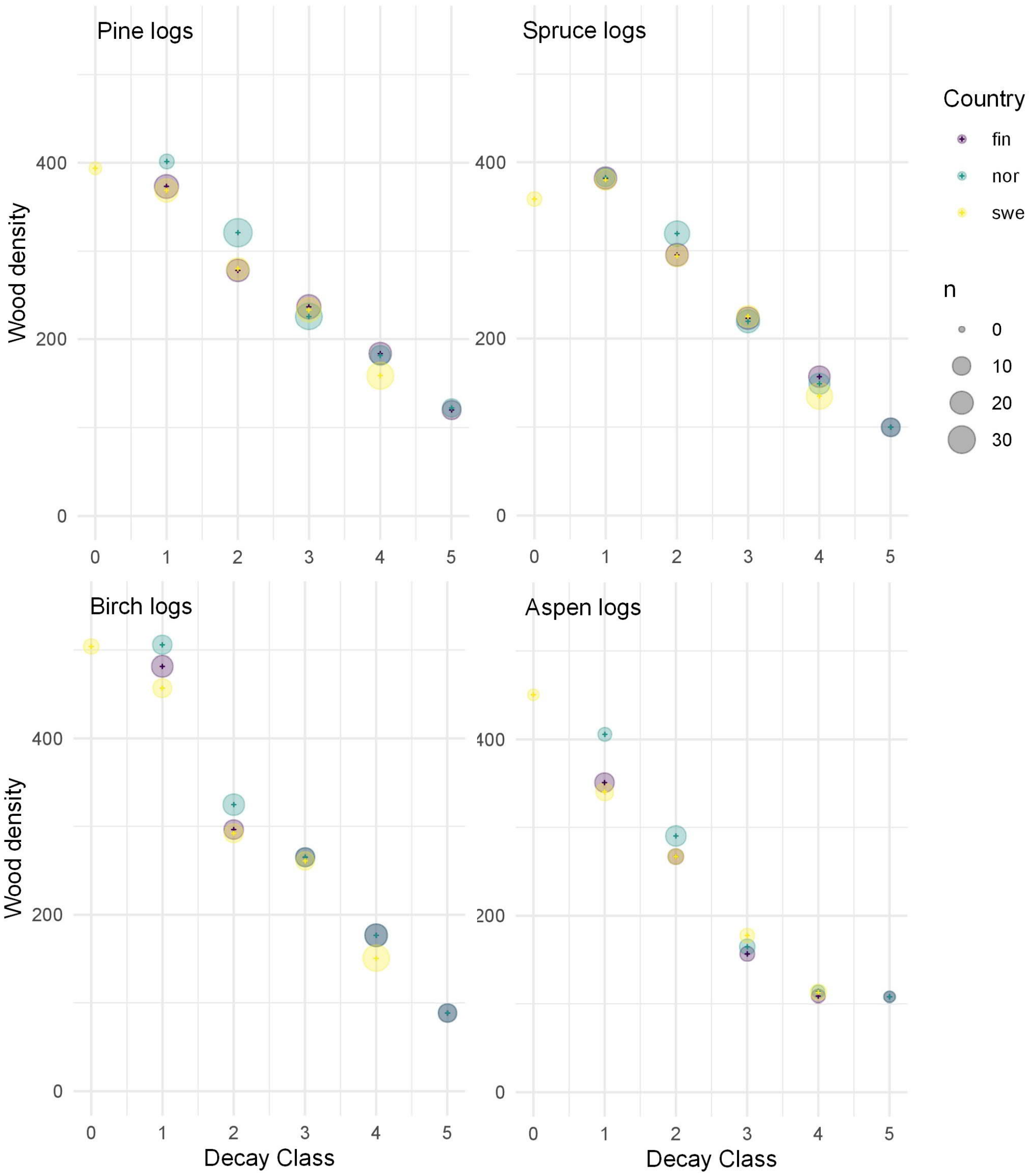


**Fig. S2b.** Average log (fallen dead wood) wood densities in the national classification systems of Finland (fin), Norway (nor), and Sweden (swe).
